# BayeSuites: An open web framework for massive Bayesian networks focused on neuroscience

**DOI:** 10.1101/2020.02.04.934174

**Authors:** Mario Michiels, Pedro Larrañaga, Concha Bielza

## Abstract

BayeSuites is the first web framework for learning, visualizing, and interpreting Bayesian networks (BNs) that can scale to tens of thousands of nodes while providing fast and friendly user experience. All the necessary features that enable this are reviewed in this paper; these features include scalability, extensibility, interoperability, ease of use, and interpretability. Scalability is the key factor in learning and processing massive networks within reasonable time; for a maintainable software open to new functionalities, extensibility and interoperability are necessary. Ease of use and interpretability are fundamental aspects of model interpretation, fairly similar to the case of the recent explainable artificial intelligence trend. We present the capabilities of our proposed framework by highlighting a real example of a BN learned from genomic data obtained from Allen Institute for Brain Science. The extensibility properties of the software are also demonstrated with the help of our BN-based probabilistic clustering implementation, together with another genomic-data example.

## 1. Introduction

Analysing neuroscience data can be particularly complex since some of these datasets can have numerous instances and/or present an extremely high dimensionality, such as fMRI or microarray data, which can be in the order of tens of thousands of variables. Learning models with massive datasets having numerous features requires unique algorithms because they may encounter the curse of dimensionality problem.

In addition to the complexity of learning in biological domains, it is particularly sensitive and risky to make decisions based on models for which the process of drawing conclusions and their implications is not understandable.

To fulfill the above-mentioned requirements, we focus on probabilistic graphical models, particularly on Bayesian networks (BNs) [1], which use probability theory to present a compact graphical representation of the joint probability distribution over a set of random variables, 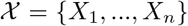. With this theoretically rich and detailed model, we require appropriate software tools to learn, visualize and interactively manipulate the resulting model, which is where the state-of-the-art BNs fail when trying to deal with massive networks.

Current state-of-the-art BN tools (e.g., shinyBN [2]) not only are lacking in proper ways to learn massive BNs but also are lacking in scalable inference and interactive visualizations. In this paper, we present BayeSuites, a new opensource framework, which is the first of its kind to overcome all of these issues in a single framework. Note also how BayeSuites is not a wrapper of existing tools into a graphical interface but is a comprehensive framework, integrating both new algorithms and existing packages adaptations to create a single tool specifically designed to fully exploit the BN’s interpretability features even for massive networks with tens of thousands of nodes. BN’s requirements are scalability, extensibility, interoperability, ease of use, and interpretability.

BNs consist of two main parts: a graph, which is a directed acyclic graph (DAG) representing the probabilistic conditional dependencies between the variables in 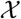, and parameters, which are a series of conditional probability distributions (CPDs) [3].

Each node in the graph represents a random variable, *X_i_*, in the vector of variables, ***X*** = (*X*_1_, …, *X_n_*), and its arcs represent the probabilistic conditional dependence relationships with respect to the other variables. Each node, *X_i_*, has an associated CPD, which represents its probability distribution conditioned on its parents, **Pa**(*X_i_*), in the graph (Figure 1). With all this information, the joint distribution of all the random variables can be expressed as

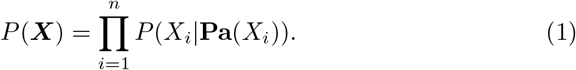

**Figure 1:**
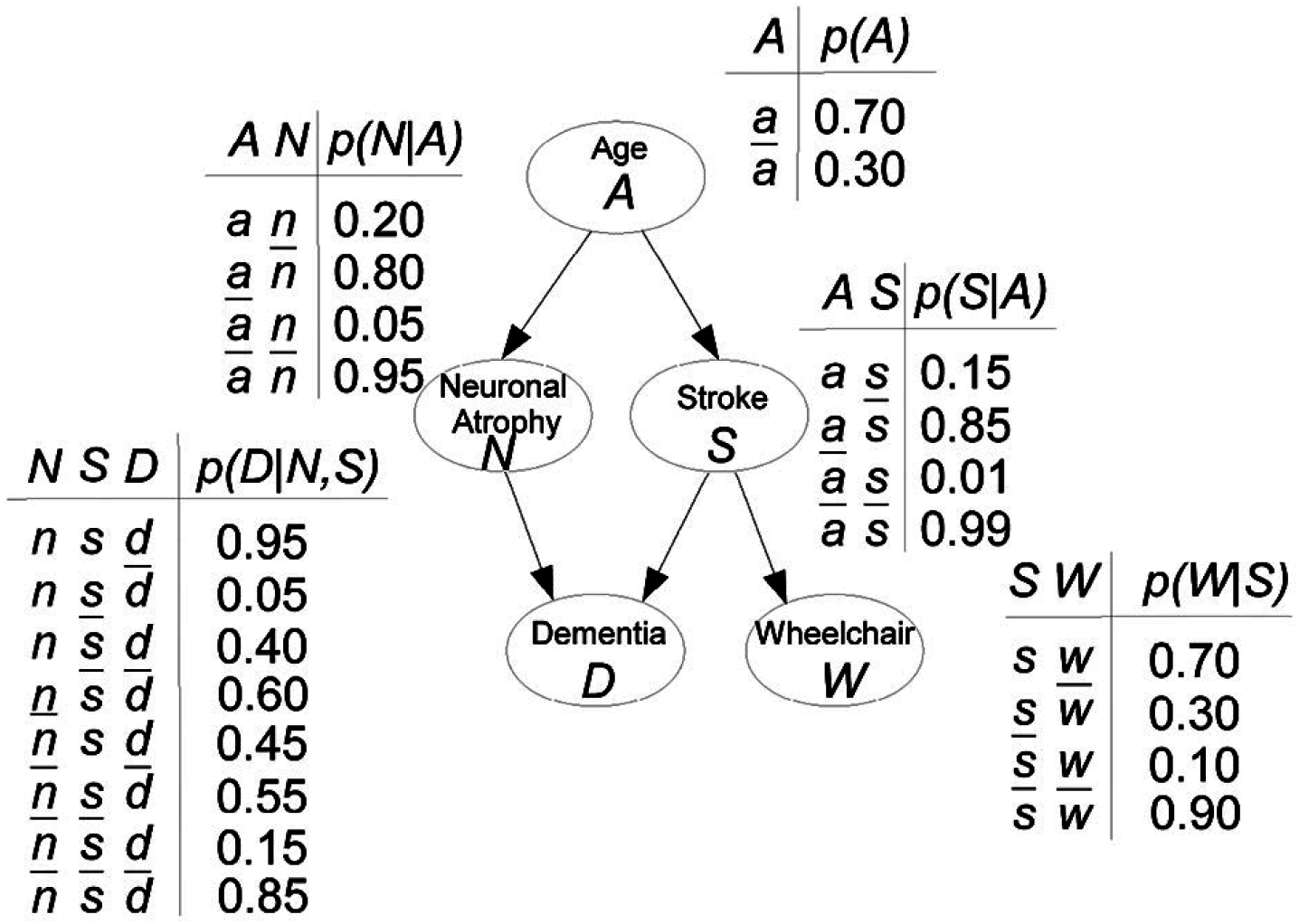
Hypothetical BN example modelling the risk of dementia. Figure extracted from [4].

The graphical component in a BN is particularly useful because it presents the probabilistic relationships between the variables. In addition, the inference machinery offers prediction and interpretability capabilities about the probabilistic reasoning and the model. For a more in-depth review of the interpretability features of BNs, we refer the reader to [5] and [6]. Owing to their interpretable nature, BNs have already been applied to neuroscience data with successful results [4, 7].

Following the example in Figure 1, if a patient has neuronal atrophy but has not had a stroke, by using inference tools, we can calculate that there is a 0.40 probability he will be demented: 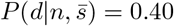.

Even if the current state-of-the-art BN tools supported massive BNs, they would not have all the proper tools for their interpretation. Visual interpretation of BNs has been studied for decades [8]. For example, [5] proposed that global network visualization should allow focus on certain parts of the structure. [9] used the arcs in a Bayesian network to show additional information; for example the thickness of an arc could represent the strength of influence between the nodes. [10] introduced a software tool providing interactive visual exploration and the comparison of conditional probability tables before and after introducing some evidence. [11] introduced multi-focus and multi-window techniques that were useful in focusing on several areas of the Bayesian network structure at the same time. Some of these advances have been implemented in major BN frameworks, which will be discussed later, but to date, there was no tool where all these features converge. BayeSuites not only focuses on scalable methods for learning and inference but also incorporates all these interpretability features with our modern implementations adapted for massive networks; in addition, it includes many newly designed methods, which are discussed in later sections.

The paper is structured as follows: In Section 2, we review the abovementioned requirements that BN software tools should meet by comparing state-of-the-art tools and highlighting where all of them lack one or more fundamental aspects so that they cannot fully express all the BN capabilities.

In Section 3, we organize the presentation of BayeSuites into the same categories as Section 2, but we explain how BayeSuites addresses all these interpretability requirements that the other BN tools failed to address in any way. We also, in this section, provide performance comparisons with other software packages, when possible. We explain the last interpretability requirement (ease of use, Section 3.4) by providing real-world use cases with genomic data. The objective of these use case examples is two-fold: (i) summarize BayeSuites capabilities in a detailed and graphical way and (ii) explain all the steps required to learn, visualize and interpret BNs with BayeSuites.

Finally, in Section 4, we discuss different use cases where the frameworks could prove useful. We also present future improvements to be implemented in this line of research. We conclude in Section 5 by providing a summary of the features that makes BayeSuites a unique framework compared to the existing BN software tools.

## 2. Problems with state-of-the-art of software in massive BN interpretability

In this section, we review the problems with the current BN software frameworks and packages by explaining the contents summarized in Table 1, which makes comparisons between the most comprehensive BN software. However, there may exist other tools of particular importance for a specific purpose, which will also be highlighted in each of these subsections below.

**Table 1:**
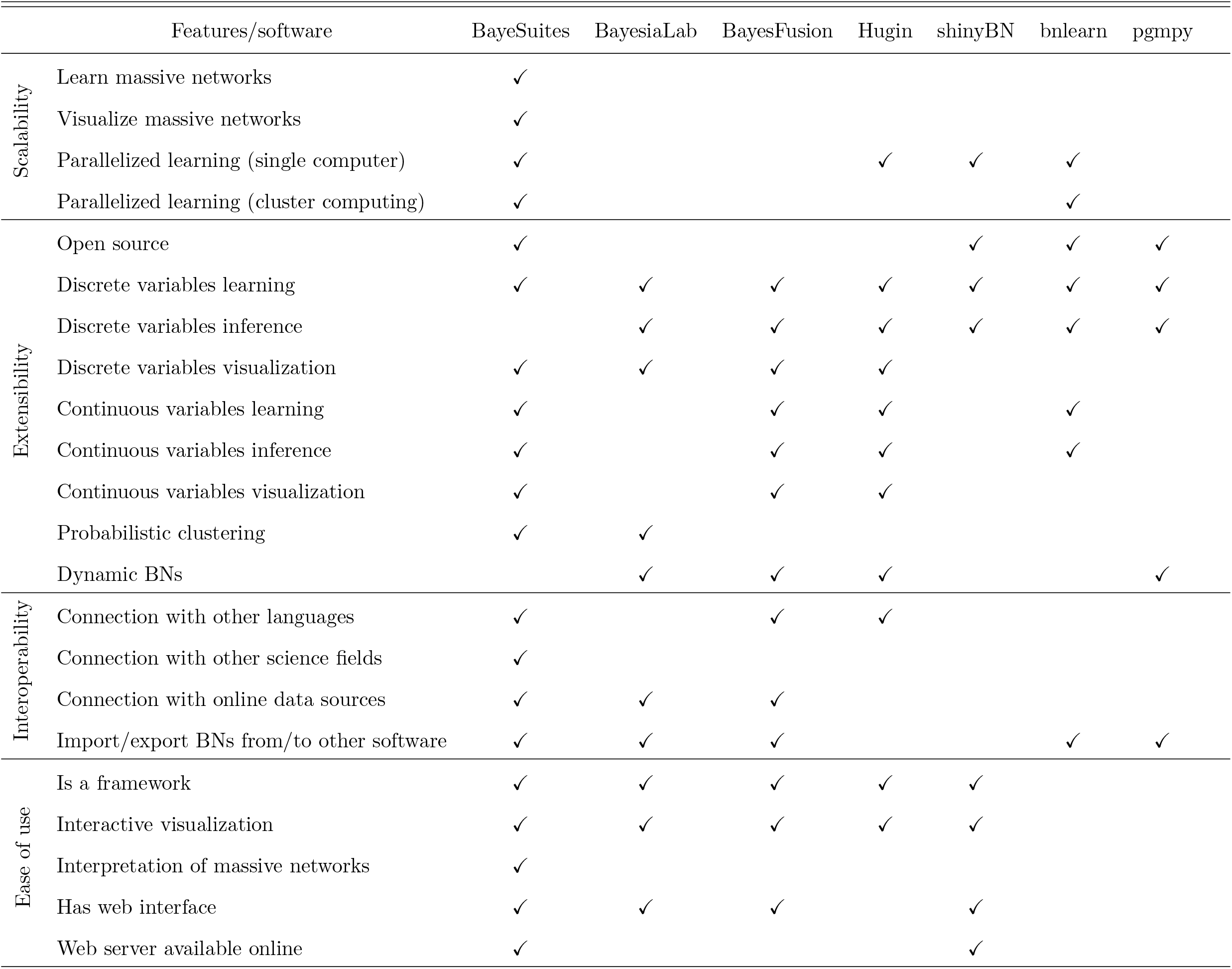
Comparison of the main BN software frameworks/packages

It is important to differentiate between individual software components addressing specific tasks (e.g. a learning algorithm), referred to as software packages, and general frameworks, as the one presented in this paper, which provide all the necessary tools to work with BN capabilities (learning, visualization, and reasoning). When we classify software as tools, we are referring to both frameworks and software package categories. Four of the major BN frameworks are BayesiaLab [12], Hugin [13], and BayesFusion [14] (which uses the SMILE engine under the hood, also with a proprietary license), and the recent shinyBN [2], which uses bnlearn under the hood. In the category of software packages, the most complete ones to date are bnlearn and pgmpy [15]. We also want to point out that we did not include other open source packages in Table 1 since most of them are outdated or nearly outdated (e.g., JavaBayes [16], BanJo [17], JNCC2 [18], BNJ [19], MSBNX [20], or Bayes Net Toolbox [21]) and/or only include very specific algorithms.

### 2.1. Scalability

Massive BNs present mainly three scalability problems: learning their graph structure, efficiently visualizing it, and developing a fast inference engine for the reasoning.

When the number of variables is extremely small, the graph structure of a BN can be even modelled with only expert knowledge. However, when the dataset has numerous random variables, the graph must be learned by computational methods. Learning the structure of a BN is known to be an NP-hard problem [22]. The search space for all the possible DAGs is super-exponential in the number of nodes, i.e., 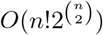 [23]. Different algorithms attempt to solve the above problem by applying heuristics to this super-exponential space.

The problem becomes comparatively more complex when dealing with a massive number of variables of the order of thousands of nodes, requiring distinct types of algorithms for constraining the computational memory and time. This problem can be solved in a reasonable time by two methods: constraining the graph structure and developing new algorithms that completely utilize parallelization technologies. The first solution includes algorithms that inherently constraint the structure (e.g., the Chow–Liu method [24]) and the generating poly-tree recovery algorithm [25]; in the latter, the resulting graph can only be a tree or a polytree. There are other algorithms which by default do not constraint the structure; however, when the problem has an extremely high dimensionality, they include assumptions, like limiting the number of parents, for each node to finish in a reasonable time. Some examples of this case are the PC algorithm [26] and the max–min hill-climbing (MMHC) algorithm [27]. These kinds of algorithms are included in most BN software tools such as bnlearn and its related packages (such as shinyBN), pgmpy, Hugin, etc. For a more detailed view of BN structure learning algorithms, we refer the reader to [28].

However, some problems like learning gene regulatory networks (GRNs) need to be modelled without restricting the structure, because all types of relations between the variables are possible. The algorithms available for these models are highly limited because most of them cannot be parallelized; therefore, new optimized algorithms are emerging [29, 30, 31]. Another problem is that some of these state-of-the-art algorithms are not typically found in the existing BN software frameworks, because the latter are not frequently updated to include new algorithms. Indeed, none of the other BN tools in Table 1 includes any of these new optimized algorithms.

Some tools such as Hugin and the bnlearn related packages support algorithms that can make use of all the CPU cores in parallel but are limited to a single CPU. However, none of the existing frameworks in Table 1 have a scalable software architecture to parallelize these algorithms on multiple computing nodes. Of the software packages, only bnlearn can run some algorithms on multiple CPUs communicating in a group (i.e. cluster computing), but these algorithms do not belong to this last category of non-restricted structure algorithms that are highly optimized for speed.

Although there exist software packages that can visualize general-purpose massive graphs such as sigmajs [32], Graphistry [33], and VivaGraphJS [34] using the GPU computational power, these are not included in any BN frameworks (Table 1). Furthermore, just including a graph visualization library with GPU rendering is not enough functionality for BNs since viewing the nodes and edges is not sufficient. We also need to visualize their node parameters and run BN operations such as making queries and computing the posterior distributions. Essentially, it is necessary to adapt existing libraries, with GPU support, to provide a rich set of interactive options to fully understand and exploit the BN graph structure and parameters. This is clearly one of the most important bottlenecks in the current frameworks when trying to deal with massive BNs since they have not even done the first step of just including a GPU library. The library would have to subsequently adapt for BNs.

Finally, we require an efficient inference engine, which in the ideal case would include exact inference. Some tools such as pgmpy, BayesiaLab, Bayes Net Toolbox, etc. include exact inference for discrete BNs, but inference in discrete BNs is NP-hard [35]; therefore, it is not a scalable solution. To reduce this cost, the network structure can be constrained with unique algorithms. This is usually the preferred option for the tools in Table 1, which sacrifices structure flexibility in favour of inference speed. An approximate inference is the alternative when we do not want to constrain the network structure; however, it is also NP-hard [36] so it is not scalable either.

In any case, most massive datasets such as fMRI or microarray data are continuous, so we need a scalable inference for continuous domains. Luckily, exact inference is tractable in the continuous space for Gaussian BNs (see Section 3.4.4). However, from all the tools compared above, only Hugin and the Bayes Net Toolbox (not in Table 1) include exact inference for Gaussian BNs. There are other tools that offer inference in continuous domains but only include it in its approximate versions (e.g. BayesFusion and bnlearn).

### 2.2. Extensibility

Extensibility refers to the software’s capability to include new functionalities easily and coherently. It is crucial for the software to be written modularly to introduce new algorithms and functionalities. Three of the major BN software frameworks are BayesiaLab, Hugin, and BayesFusion, all of which have proprietary licenses, and therefore, the code is not open-source (Table 1). This presents a significant problem in an advancing research field such as this one, because the research community cannot code its own extensions and changes. In an ideal case, the frameworks should be open-source and have simple and multiple approaches to introduce new extensions coded by the community.

Even if the commercial frameworks described above aim to be the most complete solution as possible, new extensions are limited, and this is marked by the slowness of their major updates. For example, some commercial frameworks have been developed for several years and still do not have complete support for continuous variables (e.g., Netica [37] and BayesiaLab).

In the arena of open source software, we can expect a more promising future. In the software frameworks, we find shinyBN, which incorporates a set of R tools in a coherent way, making it a good candidate for extension with new features by the software community. However, the server infrastructure is not optimal, since the shiny server is not extensible software that could be used in a large production environment.

For open source software packages, bnlearn is still the most comprehensive and widely used package for BNs. Its modular architecture and optimized algorithms have allowed it to be a robust package for more than 12 years in the software community. However, similar to other outdated packages such as Banjo [17] and JavaBayes [16], bnlearn may also be experiencing a slow decay in its extensibility features due to how much it has grown on its own. Without strong collaboration with other people, it is hard for new people to implement new features in the original C++ code since that code may not be fully documented. We also note the more recent pgmpy open source package has software extensibility that is much more attractive since it is fully coded in Python and also adheres to a good modular architecture. Indeed, its code repository is very active and there are usually new updates. We conclude that pgmpy is currently one of the best examples of software extensibility in the BN software community.

### 2.3. Interoperability

All the current frameworks in Table 1 except for shinyBN are proprietary and are specifically designed for working only with probabilistic graphical models. This lack of connection with tools from other science fields (Table 1) is a common shortcoming for both proprietary and open source tools. This means they lack connections with other statistical tools, machine learning algorithms or any other analysis and visualization tools specifically designed to overcome problems in any science field such as neuroscience, etc.

A positive feature of proprietary frameworks, as opposed to open source tools, is that they usually have API connections to other programming languages (such as BayesFusion and Hugin but not BayesiaLab) and provide input connections to general purpose online data sources (such as BayesiaLab and BayesFusion but not Hugin). However, owing to their proprietary nature, the community developers cannot implement some functionalities, such as having direct API connections with specific data sources as neuroscientific databases.

The exception to proprietary frameworks is shinyBN. Its recent appearance, however, makes it not the ideal candidate to exemplify the desired interoperability requirements since it does not have any kind of connection with other programming languages, algorithms, data sources or other BN tools (Table 1).

Finally, importing and exporting BNs from/to other tools is a fundamental feature that currently is available in almost all BN tools, but still, some tools lack it (such as Hugin and shinyBN) (Table 1).

Nevertheless, the BN community has some open-source software packages that are well maintained and have a good extensibility; however, they are designed for highly specific tasks, e.g,. learning algorithms such as sparsebn [38] and hc-ET [39] and inference packages such as gRain [40]. Other packages, such as bnlearn [41] and pgmpy [15], comprise a set of interconnected tools, but they lack some basic modules, e.g., a graphical interface or connection with other packages, which would make them considered to be frameworks. Thus, the central problem of these types of packages is the lack of completeness, unlike the proprietary options.

Furthermore, some software packages are developed for the specific purpose of a scientific research. While this is appropriate for advancing the research field, frequently these software tools are overlooked and not maintained once the corresponding paper is published. The first consequence is a waste of time associated with coding again previously written algorithms by other researchers when the existing code becomes obsolete and not extensible. Another consequence is the difficulty of integration to other software, because they may be written in a different programming language. Therefore, the library can have data format problems, software incompatibilities between versions, etc.

### 2.4. Ease of use and interpretability

Software packages regularly do not include a graphical interface; therefore, the learning curve is extremely steep for users not experts in programming, which commonly is the case with some neuroscientists. Graph visualization cannot even be considered for software packages because they mostly rely on third-party software to display a static view of the BN structure and cannot display node parameters in the graph (e.g. bnlearn, pgmpy).

In comparison, frameworks are much more user friendly because they provide a sophisticated interactive graphical interface to work with (see Table 1). However, as a direct implication of their low scalability, they are not capable of visualizing massive networks. Furthermore, even if they had the proper tools to display massive networks, none of them currently has the proper tools to interpret them (i.e. multiple specific layouts, rich filtering tools, etc). Indeed, some of them (e.g., shynyBN), do not even have a complete set of tools for interpreting small size BNs, since they lack some functionalities such as displaying BN parameters attached to each node of the graph.

Ease of use also depends on the accessibility of the tool. Web interfaces are currently robust enough to be considered as the preferred option here, since they are accessible from everywhere and are platform independent. We can see, therefore, an increasing number of tools developing web interfaces as the entry point to their software (e.g., BayesiaLab, BayesFusion and shinyBN). However, not all these tools deploy their software in their own web server to be accessible from everywhere, which forces the users to locally deploy the server in their own computers if they want to make use of their web interfaces. Indeed, only shinyBN provides its tool in an already deployed web server accessible from the Internet (see Table 1).

Moreover, in the case of proprietary frameworks, specific solutions for distinct use cases (e.g., automatically running a set of algorithms when new data emerges from a database) cannot be developed by different research teams, because of their extensibility problem. This problem is another bottleneck when customized solutions need to have an easy and rapid workflow, ranging from acquiring the data to analysing them.

## 3. BayeSuites

In this section, we present BayeSuites, whose software architecture has been specifically designed to overcome all the problems highlighted in the previous section (see also Table 1). Summarizing, we find ourselves stuck in incomplete open source solutions versus more complete solutions in the proprietary field. The objective of the framework presented in this paper is to combine the best properties of both worlds and present one complete open-source solution, with the possibility of further improvement and becoming increasingly complete.

The name of BayeSuites originates from the fact that it is embedded in the NeuroSuites platform, which we developed for integrating different neuroscience tools. Its inclusion in the NeuroSuites platform, instead of deploying it on an isolated platform, is because this tool is specifically designed to overcome large scale problems, which are common in some neuroscience topics, as in the genomics examples presented here.

BayeSuites has already been successfully used with genomic data in [31], and as genomic examples, here we present real-world use cases to illustrate how we addressed the four interpretability requirements explained above.

### 3.1. Scalability

NeuroSuites is developed as a scalable web application to run the heavy operations in the backend while providing a lightweight rapid experience in the frontend. Its framework follows a modular architecture (Figure 2), where each fundamental component is isolated as a Docker [42] microservice (Figure 2.1); therefore, the system can be easily scalable horizontally and work as a whole. Moreover, multiple monitoring tools have been included since the architecture became large and complex, and a set of tools is provided to monitor the state of the hardware, logs, task queues, etc.

**Figure 2:**
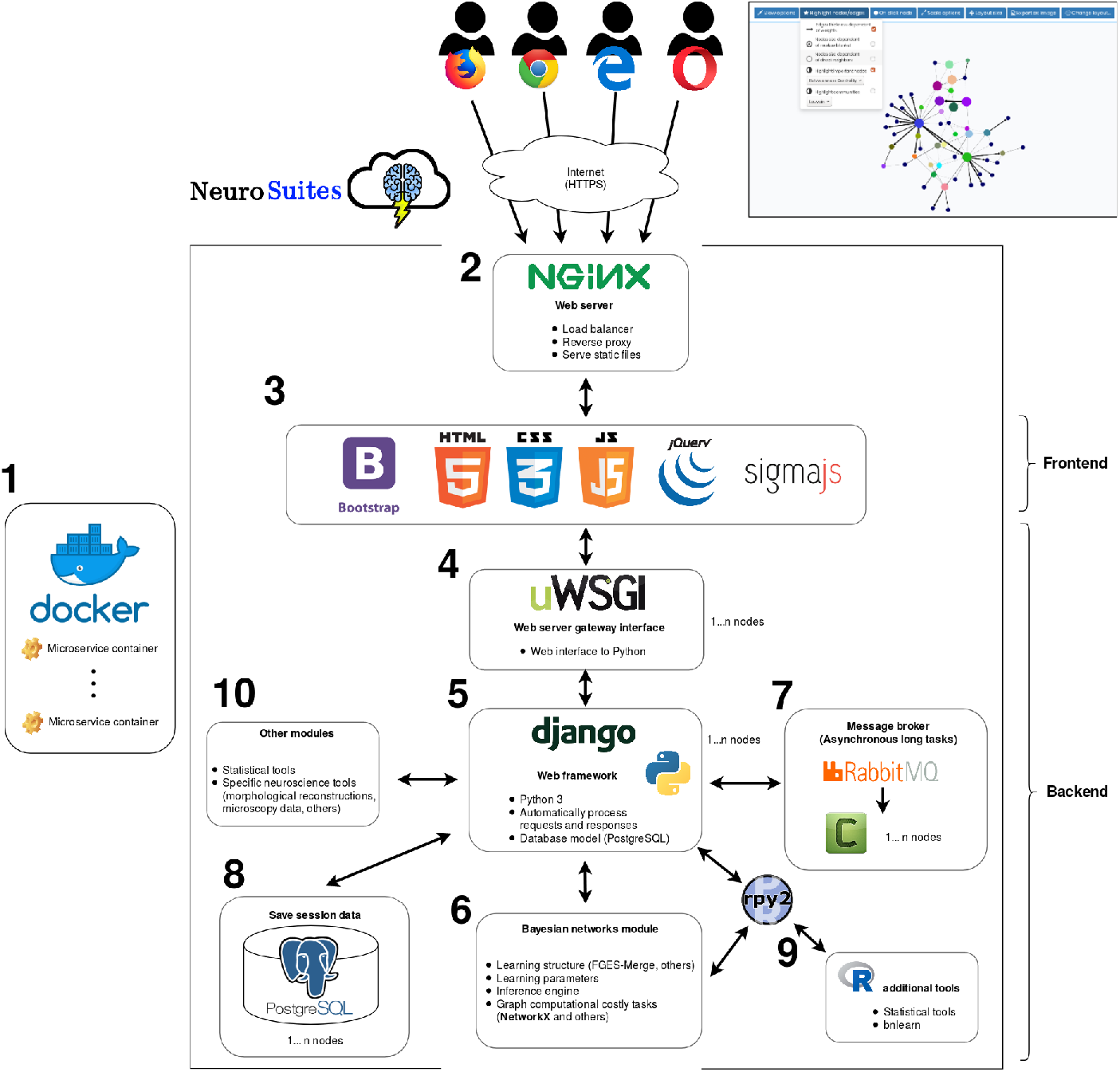
Software architecture of BayeSuites.

**Figure 3:**
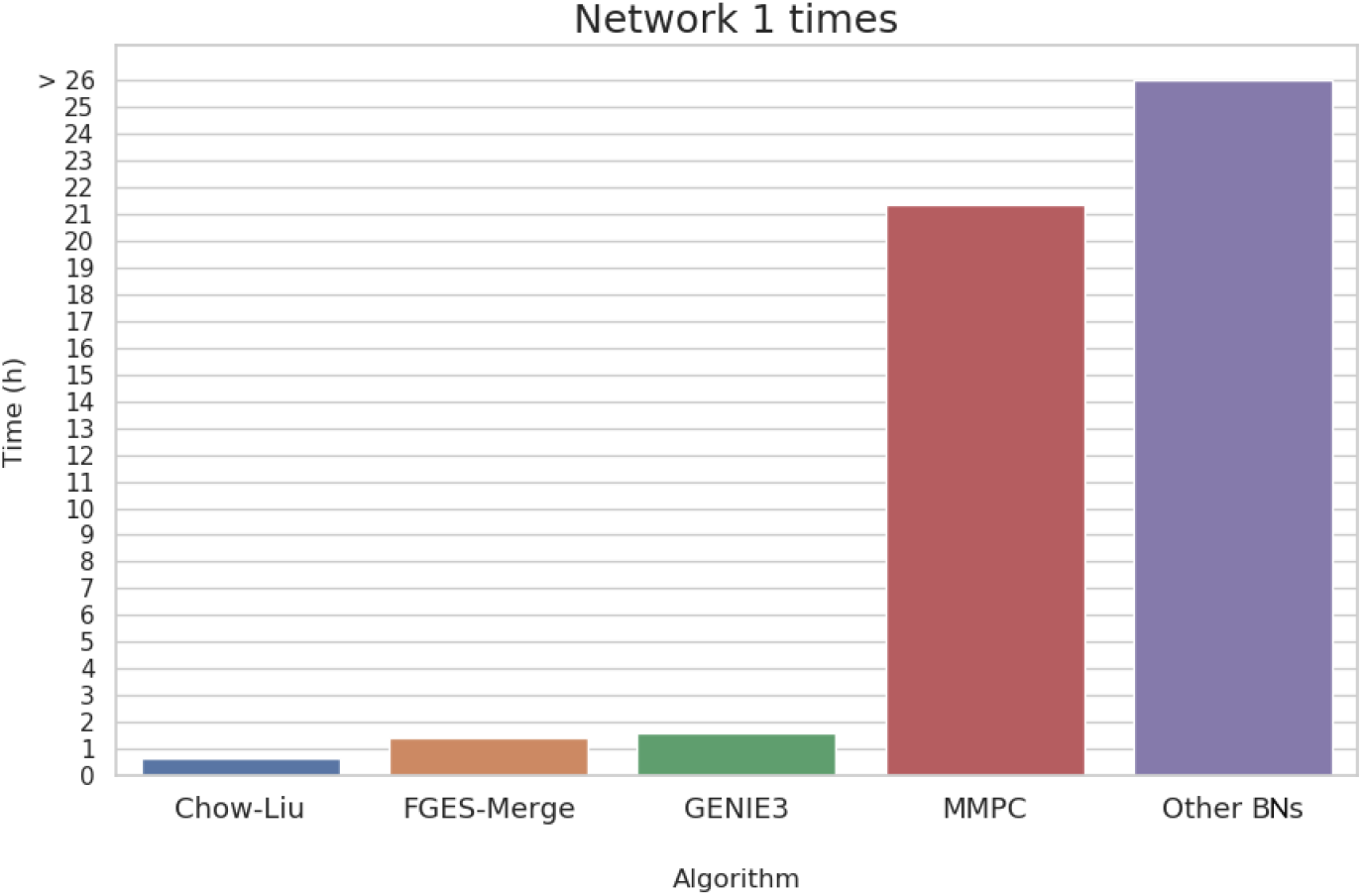
Performance time comparisons of the main BN structure algorithms. All methods were run in BayeSuites. The network learned is the Network 1 (1000 nodes) from the DREAM5 challenge dataset [57]. FGES-Merge is the method specifically designed and implemented by us for massive BNs. GENIE3 is also a method designed for large-scale networks whose original implementation was included in BayeSuites. For the other methods, their original implementations were coded in bnlearn and so were included in BayeSuites. The last category named “Other BNs”, refers to all the remaining BN methods (i.e., PC algorithm, grow shrink, hill climbing, IAMB-related algorithms, MMHC), which ran for more than 26 hours and did not finish.

The scalable architecture is designed to be efficient and solve the computational problems of visualizing and managing large learning algorithms and graph operations. The nginx web server [43] provides the entry point for the web requests (Figure 2.2) and also acts as the load balancer in case the server has multiple instances running the web application.

The frontend code (Figure 2.3) is based on vanilla JavaScript (JS) and JQuery, to provide a simple but efficient architecture, and the style is in HTML5, CSS3, and Bootstrap3. To provide a scalable visualization of the BN graphs, we have made various extensions to the sigmajs library [32], which range from visualizing the node parameters to numerous specific BN operations, fully explained in 3.4 section. Sigmajs uses a WebGL engine, which utilizes a GPU card to efficiently visualize massive graphs.

To transmit the requests and responses from the frontend to the backend, we employ the uWSGI software [44], which acts as a web server gateway interface to communicate with the Python backend (Figure 2.4). The backend core (Figure 2.5) is written in the Django framework [45], to allow us to use optimized Python libraries for managing the graph and also other scientific libraries (e.g., Numpy [46], Scipy [47], or Scikit-learn [48]) is the main library used in the backend to store the graphs and run the graph manipulation tasks. Lightweight graph operations, such as colouring groups of the nodes, are completely conducted in the frontend with the sigmajs library. The heavyweight operations are sent to the backend where they are processed with NetworkX, and the result is sent back to sigmajs to update the graph (Figure 2.6).

Standard HTTP requests and responses have time and computational limitations, which make them unfeasible to run long-duration tasks, e.g., some BN structure learning algorithms. To overcome these limitations, we have included a queue-workers system using RabbitMQ [49] and Celery [50] (Figure 2.7). The system arranges all the long time-consuming requests and queues them to be executed in the most efficient order. The system administrator can opt to scale the necessary containers when the workload is not sufficient for the number of concurrent users. For instance, when the workload is highly intense in the heavy operations, the system administrators will increase the number of workers and the queue system will automatically distribute the workload.

For high memory efficiency, the uploaded datasets are internally stored on our server using the Apache Parquet [51] format. To save the internal state of an application, the data session of the user is stored in a PostgreSQL database [52] connected to the Django framework to process all the operations in transparently (Figure 2.8).

The included BN structure learning algorithms are categorized into the following six groups: (a) Statistical based (from Scikit-learn [48], only for continuous variables): Pearson correlation, mutual information, linear regression, graphical lasso, and GENIE3 [53]; (b) Constraint based: PC, grow shrink, iamb, fast.iamb, and inter.iamb; (c) Score and search: hill climbing, hill climbing with tabu search, Chow-Liu tree, Hiton parents and children, sparsebn [38], and FGES-Merge [31]; (d) Hybrid: MMHC and MMPC; (e) Tree structure: naive Bayes, tree augmented naive Bayes; (f) Multi-dimensional Bayesian network classifier. All the algorithms where we have not specified a reference here, were implemented in bnlearn.

Only some structure learning algorithms are suitable for large-scale networks, such as the Chow–Liu algorithm, GENIE3, sparsebn, MMHC, and FGES-Merge. However, for massive networks only the FGES-Merge can learn a network in a reasonable time without constraining the structure, because it is coded to run in parallel in multiple computing instances. BayeSuites includes MPI [54], which allows this type of parallelism using the mpi4py Python package [55]. However, owing to the computational limitations of our production server, we cannot provide more than one computing node. Nevertheless, developers who install their own NeuroSuites instance can make use of this parallelization capability by deploying multiple computing nodes to run their desired Docker containers.

BN parameter learning and inference engine (Figure 2.6) have also been designed to be scalable for massive BNs and are explained in detail in Sections 3.4 and 3.4.4, respectively.

#### Performance analysis

For this performance analysis, it is important to note that BayeSuite’s goal is to be a scalable framework for massive BNs. This means our target is not to implement several learning or inference algorithms to surpass the state-of-the-art algorithms but to have a solid basis of scalable methods to be able, for the first time ever, to manage massive BNs in a user friendly interactive web environment.

#### 3.1.1. Structure learning performance analysis

Structure learning is usually the most computationally costly process when learning BNs. Comparison with other BN tools in terms of speed is not always a meaningful measure since most BN tools use the same algorithms under the hood, just with different implementations. Indeed, some frameworks such as shinyBN or even BayeSuites reuse the same implementations of bnlearn for some of their algorithms. For this reason, we compare our specific algorithm for massive BNs, which was implemented by us, named FGES-Merge [31], with the most common structure learning algorithms implemented in bnlearn. Moreover, we include the GENIE3 algorithm in the comparisons since it is specifically created for large scale problems such as gene regulatory networks (GRNs). The network learned in this test is the Network 1 (1000 nodes) from the DREAM5 challenge dataset [56]. We ran the test on this medium network to be able to compare our algorithm with others. Running the same test with a larger network with 20,000 nodes would not be possible since other algorithms would run for a very long time without finishing. This test shows not only that FGES-Merge improves the speed in comparison to other algorithms (except for the Chow-Liu algorithm, which is expected since it limits the graph structure to be a tree) but also that algorithms can run for a long time in BayeSuites without any network/memory problems thanks to its scalable architecture with asynchronous tasks (see Figure 2.7). It is important also to acknowledge that FGES-Merge is implemented with parallelization capabilities and therefore is executed in 3 computer nodes, while the other algorithms do not support this parallelization, so they are run on one computer node. Every computing node ran on Ubuntu 16.04, Intel i7-7700K CPU 4 cores at 4.2 GHz, and 64GB RAM. In terms of structure recovery benchmarks, it was proven that FGES-Merge outperforms existing BNs methods for the largest GRN of the DREAM5 challenge dataset, corresponding to the *Saccharomyces cerevisiae* network (5,667 nodes) (see [31] for a detailed comparison).

#### 3.1.2. Inference performance analysis

As reviewed in Section 2.1, inference performance is critical, even for medium size networks. BayeSuites has implemented exact inference for Gaussian BNs (see Section 3.4.4), which makes it possible to resolve inference questions in < 5s for small networks (approximately < 300 nodes and edges). The interesting point here is the scalability nature of this algorithm, which makes it possible to run inference for massive networks in less than 30-40 seconds, even in the network in the example of Section 3.4.3, that has 20,787 nodes and 22,639 edges. It is also critical to note that the resulting multivariate joint distribution and the original joint distribution are cached in the backend. This means that this 40-50 s process is done only when evidence or a group of evidence is fixed. Once this is done, any query operation with these parameters are nearly instantaneous (< 2 s).

Again, here, it is not possible to make meaningful comparisons with other BN frameworks because they do not include exact inference for Gaussian BNs, except in the case of Hugin. However, Hugin does not support deploying massive networks, so it is complicated to run these tests, although we assume that performance should be similar to ours if massive networks were supported. To just get a grasp of the times for running inference for other algorithms applicable to Gaussian BNs, we can see in [57] how typical sampling algorithms such as Gibbs sampling takes more than 8 minutes to run inference in the ANDES network (223 nodes and 338 edges). However, also in [57] we can see how new algorithms are clearly outperforming these older sampling algorithms, such as new importance sampling algorithms that can run inference in 8 s for the same ANDES networks. All these sampling experiments were run in SMILE (the computational engine of BayesFusion). Another promising research line is variational inference, which shows a good performance time of around 9 s for a randomly created network of 500 nodes and 1000 edges [58]. In summary, all these advances perform well for medium size networks, even for other parameters different than Gaussian distributions, but for now they cannot reach the performance of exact Gaussian BNs in terms of accuracy and speed.

#### 3.1.3. Visualization performance analysis

Performance times to load small networks is similar to other BN frameworks (i.e. < 2 s for networks of approximately < 500 nodes. For massive BNs this time is increased (about 10-15s for networks of approximately 20,000 nodes and 20,000 edges). However, performance comparisons with other BN frameworks for massive BNs is not possible since they do not even support the visualization of these networks. Hence, when trying to load massive networks, any kind of computational problem can arise, but mainly graphic problems strike since these frameworks do not use GPUs. Moreover, even if they supported massive networks visualization, they are not prepared to properly manage them with multiple layouts and filtering tools. These BayeSuites functionalities are discussed in detail in Section 3.4.1. To advance, running one of the proper layout algorithms for massive BNs takes less than a minute (or even less since some are iterative algorithms that can be stopped at any moment) to provide a clear and coherent graph visualization. They run without needing computational power on the user’s computer since most of these layout algorithms are executed in the backend.

Comparison of two BNs superposed in the same window is another unique functionality of BayeSuites (Section 3.4.3), which is highly optimized since actually only one graph is maintained in memory, but edges are specific to either one or both BNs. The performance speed, therefore, is instantaneous when changing between the two BNs once they have been loaded. But again, performance speed comparisons with other BN frameworks is not possible since they do not provide this kind of BN comparison.

#### 3.1.4. Server performance analysis

Finally, server performance is also an important metric in the case of web architectures. As a performance example, we can see an uptime of 99.8 % for the last month thanks to the extensive use of monitoring tools for the server deployment. Downtimes were only caused by necessary updates, which are fast and nearly automated. Moreover, periodic backups ran during non-excessive use hours such as nighttimes to improve the performance of the server during working hours.

### 3.2. Extensibility

NeuroSuites follows an extensible architecture where each module has internal submodules, allowing the modules to extend in any manner.

This extensibility enables highly easy integration of new learning algorithms or new graph functionalities for BNs and other modules. For example, to include a new structure learning algorithm, the only requirements are taking the dataset as a Pandas data frame [59] and outputting a NetworkX graph object. The changes in the frontend would be minimal, only adding a new button to run the new learning algorithm. The entire workflow is automated, and the learning algorithm would be directly queued to the system when a request is sent.

As the backend core is written in Python, the easiest method to extend it is by coding a Python extension. Because we aimed to support maximal scientific communities, we also included bindings to the R programming language for the BN learning algorithms and other statistical packages. The binding was easily achieved via the wrappers provided using the Rpy2 package [60] (Figure 2.9).

To demonstrate the extensibility of the models, we also included support for BNs-based clustering models. Thus, in the backend side, a subclass of the BN model was created with the necessary extensions, and for the frontend side, the same Javascript core for BNs was recycled and the necessary extensions were included (see Section 3.4.4).

### 3.3. Interoperability

To provide an interoperable ecosystem, we designed a well-defined workflow consisting of first uploading the raw dataset and then selecting the desired tools to analyse it. Therefore, different sections can be found on NeuroSuites, where each refers to a tool or a specific set of processing tools. The focus of this study is on the BNs section; however, users can also use other tools already integrated in NeuroSuites. Some of these tools, such as the statistical analysis Section (Figure 2.10), can provide significant preliminary analysis for improved better understanding of the data to then create better BN models.

As a use case regarding interoperability, there exists an API client that can connect a data source; it is the latest live version of the NeuroMorpho.org database. This type of direct connection to a data source is convenient when the data from a specific neuroscience field are required to be connected. This allows the users to easily access the data without the need to first download the data on their computer and then upload them to the NeuroSuites server. Thanks to the extensibility properties of NeuroSuites, it would be straightforward to implement numerous data source connectors to any database, e.g., the Allen Cell Types Database [61] and the DisGeNET database for genes-human disease associations [62].

### 3.4. Ease of use and interpretability

Here, we review the capabilities of BayeSuites by presenting a complete real use case: learning and interpreting the GRN of the full human genome using brain data extracted from microarrays, provided by the Allen Brain Atlas [63]. The dataset consists of 20,708 protein-coding genes as predictor features with 3500 samples; therefore, each element in the dataset corresponds to a measurement of a gene expression level.

In step 1, the desired dataset (Figure 4a) is uploaded. In our deployed production server, we accept CSV and Apache Parquet gzip formats. Note that the BNs can also be created by different software, e.g., BayesiaLab or bnlearn, and then be imported in a BIF/CSV/Apache Parquet format to BayeSuites to visualize and interpret the model. However, for this example, we present the entire workflow to create and interpret a new model.

**Figure 4:**
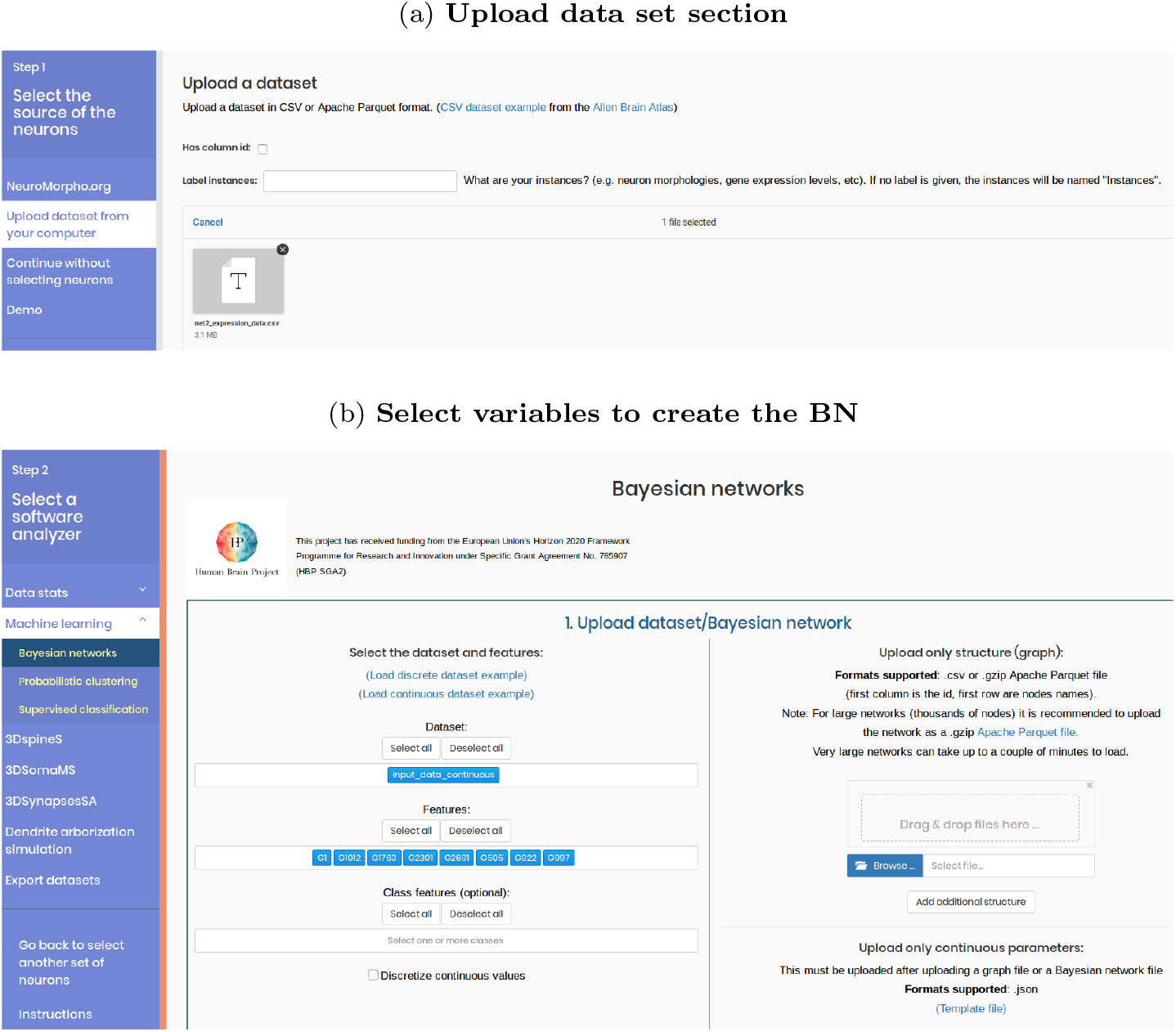
Steps to upload a data set and select its desired variables.

In step 2, we move to the BNs section under “Machine Learning” and select the desired variables to learn the model (Figure 4b). For this example, we select some continuous variables that correspond to the expression level of some genes. It is also possible to discretize the selected variables with different methods or select the class variables for a supervised classification model; however, this is not the case in our example.

Following the selection of the desired variables, the BN structure graph is learned by selecting a structure learning algorithm, as described in the field below (Figure 5a). For this example, we use FGES-Merge because it is specifically designed for genomic data, being memory and computationally efficient and having the ability to readjust the final structure to follow the topology of the GRN [64].

**Figure 5:**
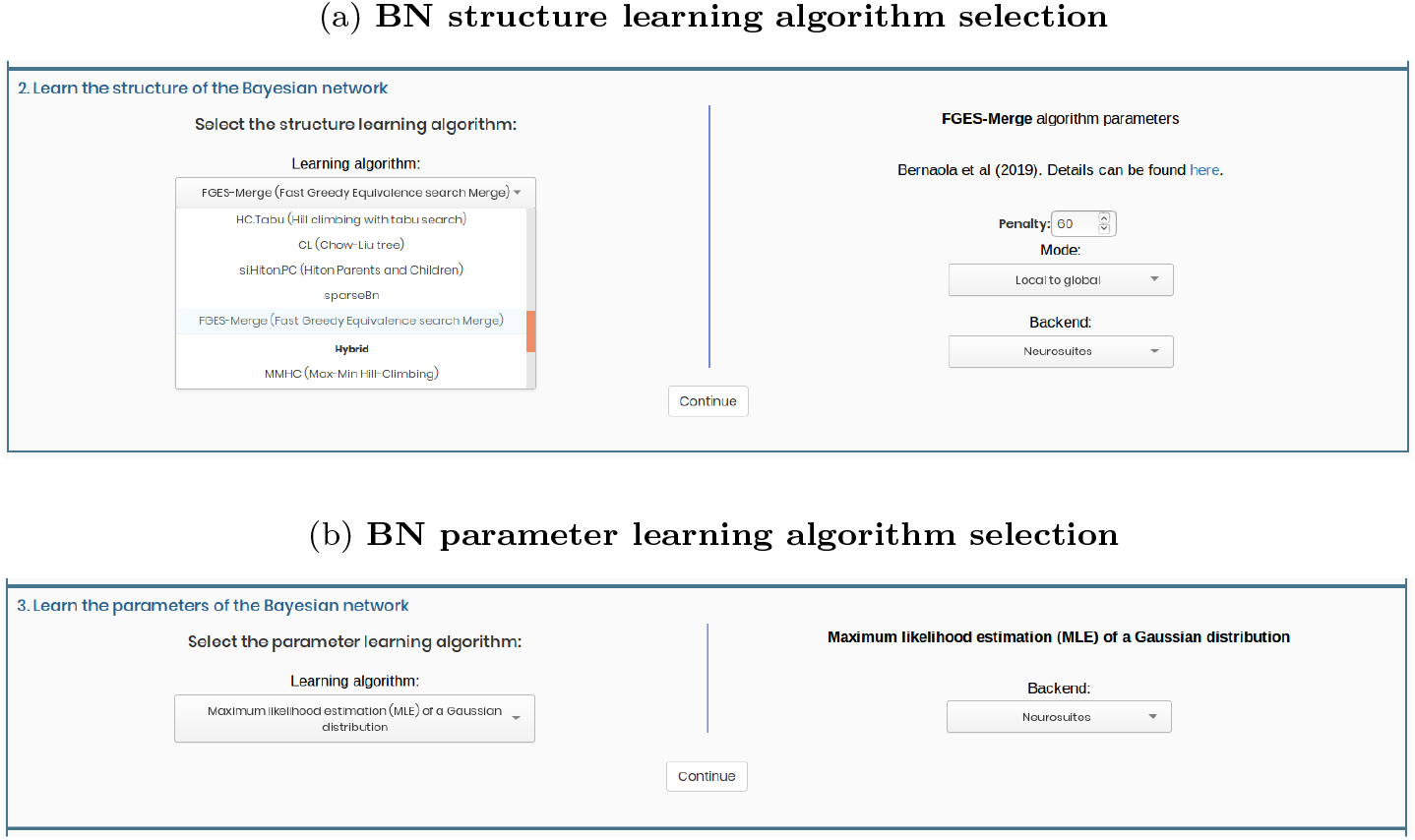
Steps to learn a BN.

Once the algorithm is completed, the obtained graph is displayed in the visualizer, and we can immediately manipulate it. Nevertheless, to provide a complete example, we also present how to learn the model parameters for each node. For this, we select the maximum likelihood estimation (MLE) of a Gaussian distribution (Figure 5b), which provide the learned Gaussian distribution for each node and the weights corresponding to the relationships with its parents. Mathematically, the CPD of a node, *Y*, given its parents **Pa**(*Y*) is

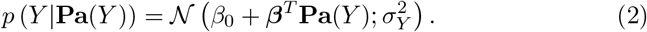

To estimate the parameters, *β*_0_, ***β***, and 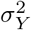, for each node, the Gaussian MLE learns a multilinear regression between *Y* and **Pa**(*Y*). The regression coefficients provide estimations of *β*_0_ and ***β***, and the mean of the regression residuals sum of the squares yields the 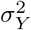 estimate.

Having learned the node parameters, we can utilize the inference engine by asking some queries to the BN and obtain the predicted results when some node values are fixed, as explained in detail in Section 3.4.4.

There are several visualization and interpretability options, which are categorized into four groups: layouts, general viewing options, highlighting nodes/edges, and parameter visualization and inference.

#### 3.4.1. Layouts

A predefined layout is displayed in the visualizer when the BN is loaded for the first time, but depending on the problem, a different one might be needed to be set. Choosing the appropriate layout should be the first step to understand the graph structure of a BN. The layouts (Figure 6a, right corner) can be tree-based layouts (Dot [65], Sugiyama [66]), force-directed layouts (Fruchterman-Reingold [67, 68], ForceAtlas2 [69, 70]), circular, grid, and image layouts. The last one is a novel method developed by us to create a layout by detecting the edges from any image. It is particularly useful for creating user-defined layouts or complex layouts that cannot be implemented by other algorithms. Layouts are computed in the backend side for efficiency, although we also provide a frontend (client version) implementation for the ForceAtlas2 algorithm [71].

**Figure 6:**
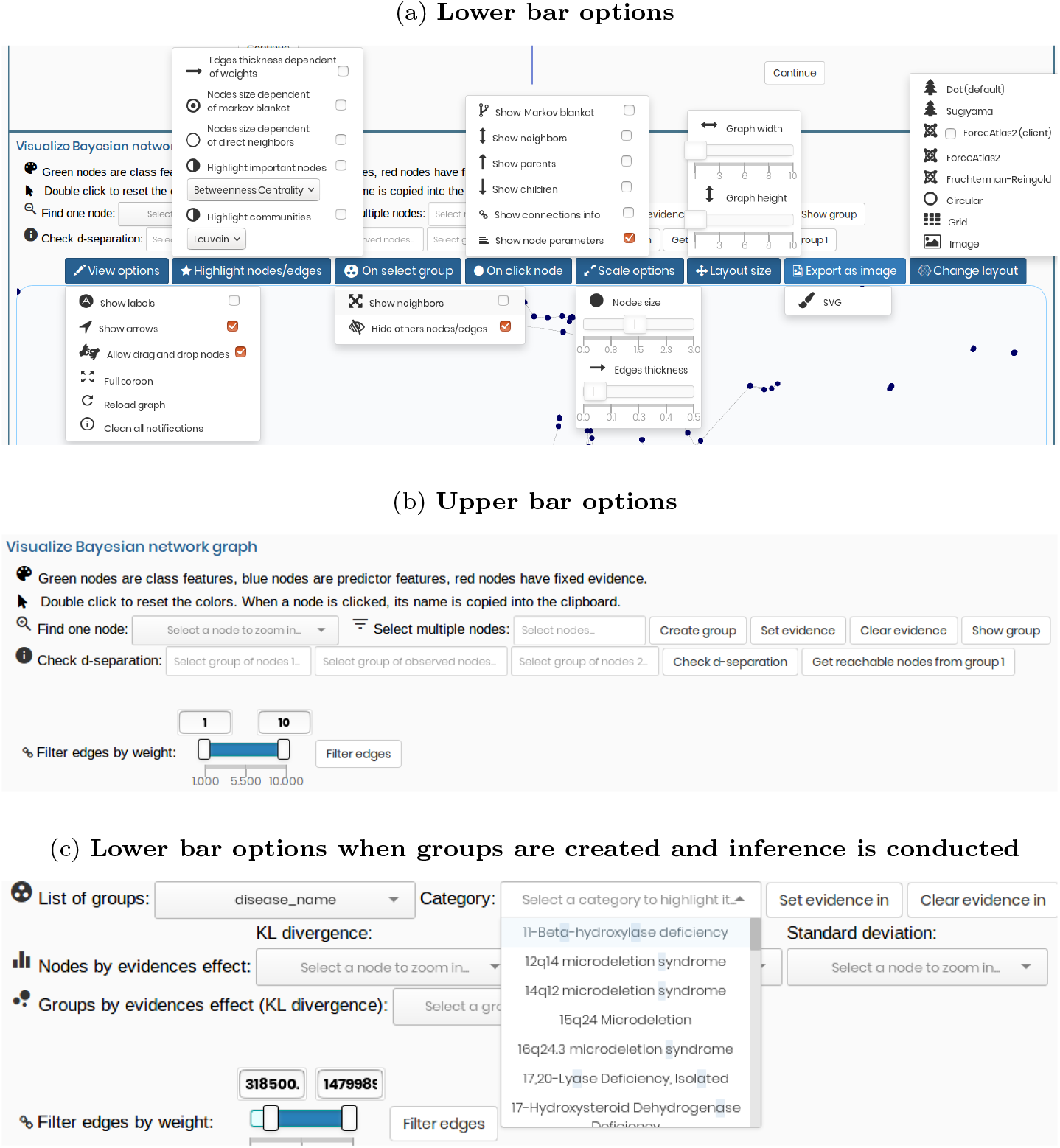
BN visualization options.

For small or medium BNs, tree layouts are recommended, whereas force-directed layouts are recommended for large BNs, because with this type of layout cluster formation occurs. In this example, we select the ForceAtlas2 algorithm because it can clearly yield the topology properties of GRNs (locally dense but globally sparse) (Figure 7a). Note that the extensibility nature of a project affect the convenience for the developers to add new layout algorithms or modify the existing ones to meet their own needs.

**Figure 7:**
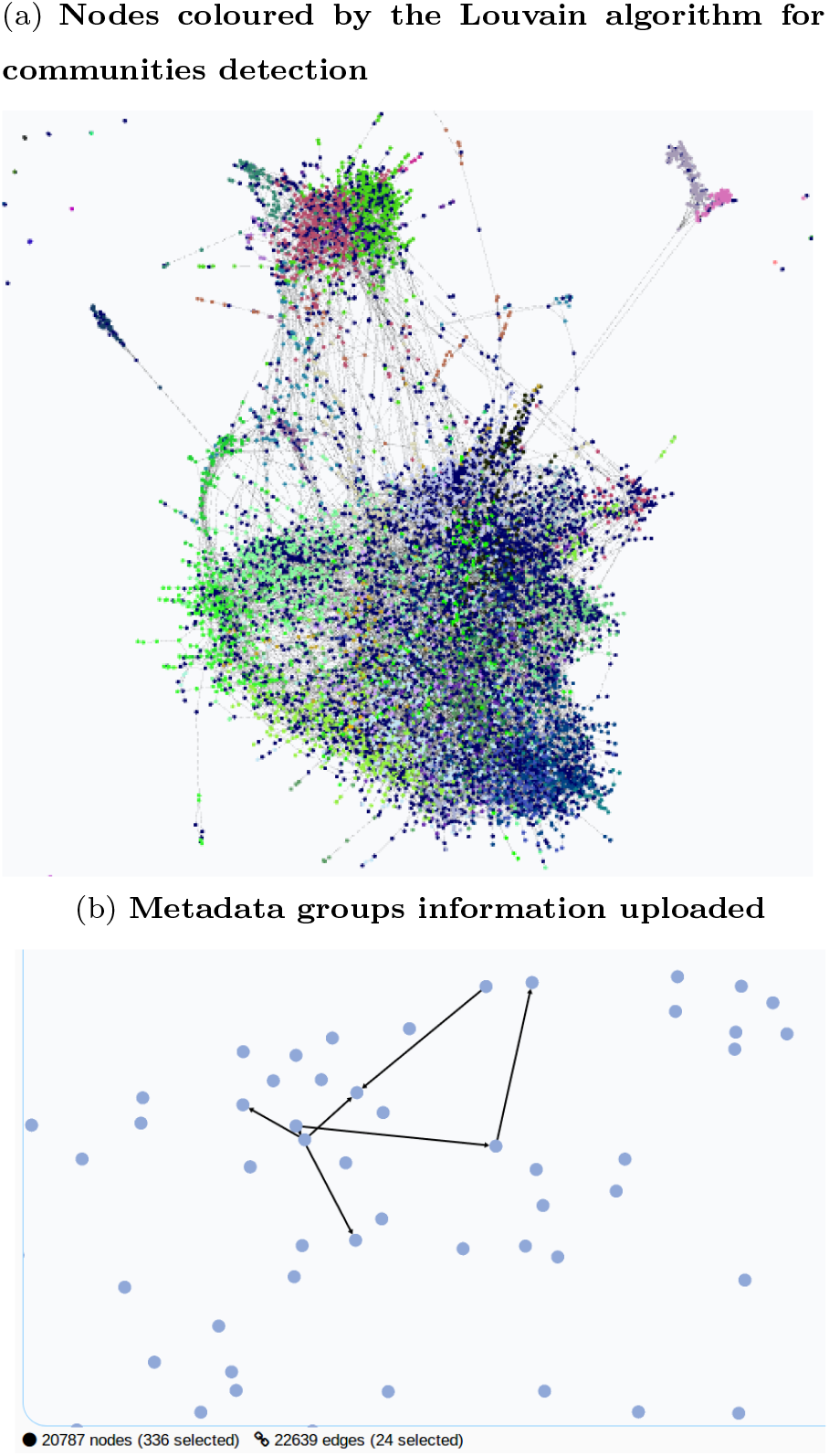
BN structure of the full human brain genome, where independent nodes are not shown. (a) ForceAtlas2 layout is applied. (b) Same network as in (a) but now only a subset of the nodes associated with the schizophrenia disease and the edges between them are selected.

#### 3.4.2. General viewing options

For general viewing options, we can easily navigate through the graph, allowing to zoom any region of interest. The lower bar of the visualizer has buttons to show/hide the labels for each node, arrows, drag and drop nodes, full screen, and reloading the graph (Figure 6a, left side).

Multiple relevant scale options are also implemented (Figure 6a, right side), such as node sizes dependent on the number of nodes in their Markov blanket or edge thickness dependent on their weights, irrespective of their reference. For instance, the edge weights can correspond to a score that refers to their importance in the network, such as the BIC score [72]. It is a penalized likelihood of the dataset calculated with the BIC difference of adding that edge versus not adding it. A filtering option to remove the edges below or above a certain weight threshold is also included (Figure 6b, bottom left).

#### 3.4.3. Highlighting nodes/edges

Subsequent to selecting the appropriate layout and configuring the general viewing options, the next step is highlighting the relevant nodes or edges. We provide tools for highlighting the nodes isolated in the Markov blanket of a given node or its parents or children (Figure 6a, centre).

When dealing with massive networks, one of the most important features is the creation of groups. The groups can be created by three ways: manually, automatically, or uploading a list of already defined groups of nodes. A node or a set of nodes manually can be selected by searching for them by their name in the search fields with auto-completion (Figure 6b, middle left, “Find one node”). Once we have selected the desired nodes to highlight, we can opt to create a group with them, and our node selection is saved to be used subsequently (Figure 6b, upper middle, “Select multiple nodes”). A name and colour can also be assigned to each created custom group.

To generate groups automatically, we can run some algorithms designed for community detection, such as the Louvain algorithm [73], which optimizes a modularity score for each community. In this case, the modularity evaluates the density of edges inside a community compared to that of the edges outside it. To select groups already created externally, we can upload the metadata JSON file, so that each node has some associated tags.

Finally, we can select a specific group (Figure 6c, upper left), and each node is displayed according to the colour of its category (Figure 7a). Moreover, we can select a specific category within a group (Figure 6c, centre), and only the nodes with that category are shown (Figure 7b).

When selecting a group of nodes, the arcs between these nodes are also selected to provide a clear view of the group. A user can also opt to highlight the neighbours of the nodes for that group, even if they do not belong to that group (Figure 6a, centre). Finally, to realize a clear understanding of where a group is within the global network, a user can enable an almost transparent view of all the other nodes that are not in the selected group.

Additionally, individual important nodes can also be selected by fixing a threshold for their minimum number of neighbours. An automatic approach has also been included to highlight the important nodes using the betweenness centrality algorithm (Figure 10a) implementation in NetworkX. It can detect the importance of a node is according to the number of shortest paths (for every pair of nodes) that pass through the node.

Comparisons of two different BNs are also possible by displaying both structures in the same graph and colouring the edges depending on which network they belong to. To achieve this, we must first upload a BN or learn it from a dataset, and then repeat this with the second BN. However, a visual comparison is not sufficient when the networks are large. Hence, we include a summary table displaying some structural measures, such as the accuracy, precision, recall, F-Score, and Matthews correlation coefficient, which use the confusion matrix of the edge differences of the second BN with respect to the first BN.

#### 3.4.4. Parameter visualization and inference

The next step is to visualize the node parameters and make some queries to the BN, to demonstrate how the inference engine works. BayeSuites supports visualization for both discrete and continuous nodes. In the case of discrete nodes, the marginal CPD table is provided, whereas in the continuous case, an interactive plot of its marginal Gaussian PDF is displayed (Figure 8a).

**Figure 8:**
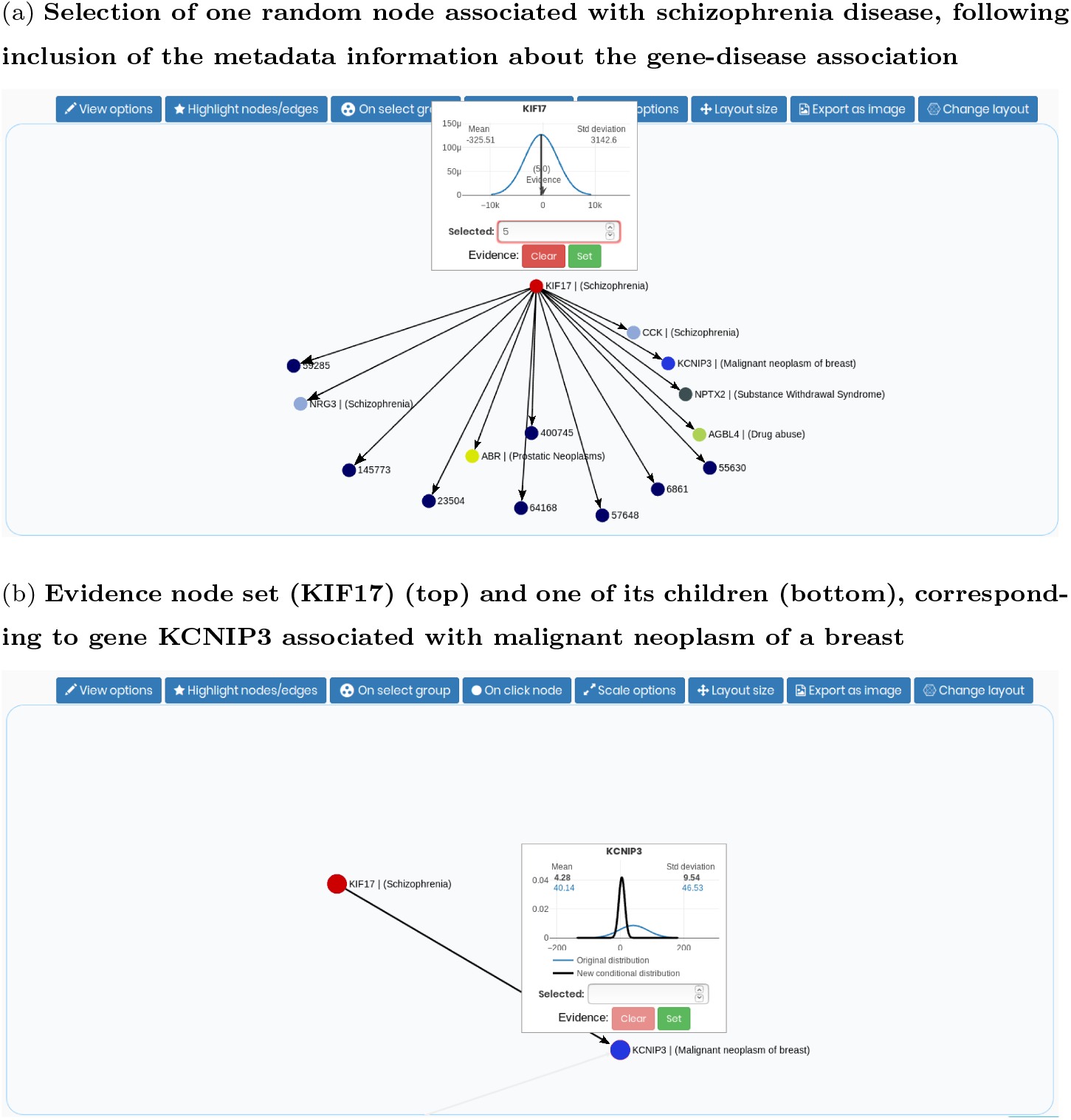
Inference workflow in BNs. The network corresponds to the full human brain genome from the Allen Institute for Brain Science. (a) In this case, we select the node on top, corresponding to gene KIF17, fix its value to make it an evidence node, *E* = e, and only show its children to have a clear view of their relations. (b) The plot includes its original marginal Gaussian PDF in blue, *p*(*Q*), as it is before setting any evidence, and the new one in black, *p*(*Q|E*), which corresponds to its PDF after setting the evidence of gene KIF17. The exact parameters are also displayed. Therefore, the inference process demonstrates how fixing a low value for the gene associated with schizophrenia (KIF17) also results in a value near zero for the gene associated to the malignant breast neoplasm (KCNIP3), which indicates a relationship between these two genes.

Because our example has only continuous Gaussian nodes, we describe the continuous exact inference engine. This involves converting the network parameters into a multivariate Gaussian distribution, 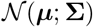; therefore, the marginalization operation for a query variable, *p*(*Q = q*), is trivial, because we only need to extract its mean and variance from the multivariate Gaussian distribution. For the conditioning probability density of a query variable given a set of evidence variables, *p*(*Q = q*|***E***), we also utilize the multivariate Gaussian distribution, following the equations described in [3].

Performing the inference operations in this manner allows a highly rapid inference engine because the most time consuming operation is conditioning over a large set of evidence variables in ***E***, which is *O*(*l*^3^), being *l* is the number of evidence variables ***E*** to condition on. This complexity is directly a result of the formulas for conditioning, as it is needed to invert a matrix of size *l* × *l*.

From the user perspective, this entire process is transparent, which is a key factor for the ease of use and interpretability of BNs. The inference process is as follows: to set the evidence nodes, ***E***, the user either clicks on the desired node and fixes the exact value (Figure 8a) or selects a group of nodes. The last option only allows fixing a shared value of the evidence for the whole group, because the standard deviation of each member of the group varies from its mean value. Setting different values at each node would be inefficient because the group can be large and the nodes can have different scales.

To view how the parameters of the query nodes, *p*(*Q* = *q*|***E***), change, the user clicks on a specific node and both the original and new distributions are shown in the same plot, allowing a better comparison of how the parameters changed (Figure 8b). Note that when no evidences are selected, only the original marginal distribution, *p*(*Q = q*), is displayed on clicking or searching a desired node in the search bar. As both the original and updated distributions are cached in the backend, the estimated time for presenting the marginal distribution of a specific node is highly optimized having a constant complexity, which in real time is equivalent to only a couple of seconds.

To provide useful insights about the inference effects, we display multiple sorted lists of the query nodes, demonstrating how much their distribution changes according to the KL divergence value, mean variation, and standard deviation variation (Figure 9). When the case groups are created, a list of the multivariate KL divergence values for each group is also be displayed.

**Figure 9:**
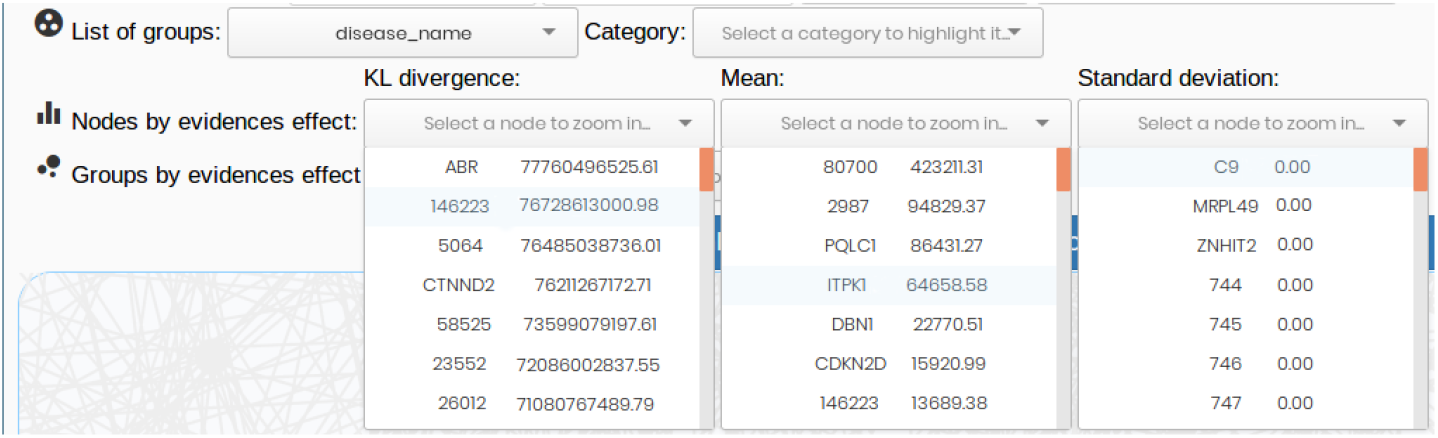
Inference effect in the query nodes. We can now infer the extent the evidence of a node (or group of nodes) affects the PDF of other nodes or group of nodes, *p*(*Q|E*), by examining the Kullback–Leibler (KL) divergence between the original and the posterior distributions or their mean or standard deviation variation. The left column in each dropdown box corresponds to the genes id, and the right column presents the score values. Note that in this example, the standard deviation values seem to be zero, because they are rounded to two decimals. Further, the effect of fixing the evidence of only one node in a network of more than 20,000 nodes can be minimal for the standard deviation of the other nodes.

In addition, to support another functionality for understanding the graph, we implemented the D-separation criterion following the reachable algorithm described in [3], which can automatically check for conditional independences. Two random variables *X* and *Y* are conditionally independent given a random variable *Z*, if for any assignment of values *X = x, Y = y, Z = z*, knowing the value of *X* does not affect the probability of *Y* when the value of *Z* is already known, i.e., *P*(*Y|X, Z*) = *P*(*Y|Z*). Thus, the D-separation algorithm can be particularly useful when we are running inference and want to determine whether some nodes are conditionally independent when some evidence nodes are given.

We have implemented further extensions to support BN-based probabilistic clustering models. The utilized dataset for this use case is also from the Allen Brain Atlas, specifically the one in the Cell Types Database: RNA-Seq Data, which contains single-cell gene expression levels across the human cortex. Therefore, the genes correspond to a set of continuous attributes ***X*** = {*X*_1_, …, *X_n_*} for the cell measurements (i.e., the dataset instances).

In model-based clustering [74], it is assumed that the data follow a joint probability distribution, *P*(***X***), which can be expressed as a finite mixture of *K* components. This implies that each mixture component, *P*(***X**|C = c*), refers to the CPD of ***X*** variables given a cluster, *c*, where the hidden cluster, *c*, has its own probability distribution, *P*(*C = c*). Thus,

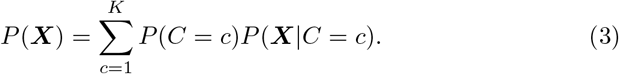

Learning the parameters for this mixture model requires a more advanced technique than MLE, because the cluster variable is hidden. Therefore, we learn the parameters (mixture weights *P*(*C = c*) and the CPD parameters, i.e., *P*(***X**|C = c*)) with the expectation maximization algorithm [75] because it can handle incomplete data.

In genomics it is typically assumed that *P*(***X**|C = c*) follows a multivariate Gaussian distribution, 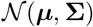. Hence, the parameters are the mixture weight vector, ***π***, and the multivariate Gaussian distribution parameters, i.e., the mean vector ***μ***, and the covariance matrix, **Σ**.

Numerous genes require a high-dimensional model, which can lead to major computational problems, in terms of both memory and time. For instance, we would have to work with **Σ**, which is an *n × n* matrix, where *n* is the number of ***X*** variables (genes in this case). To reduce the computational complexity and improve the interpretability, we can factorize this expression to encode the conditional independences between the ***X*** variables in a cluster. This allows different graphical models for different clusters, because the relationships between the ***X*** variables are conditioned on each cluster as

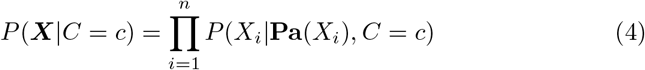

To represent this, we display each graph corresponding to *P*(***X**|C = c*) in the same BN, colouring the edges with different colours for each cluster (Figure 10a). Selection tools are also implemented to show/hide the different cluster edges and filter them (Figure 10b).

**Figure 10:**
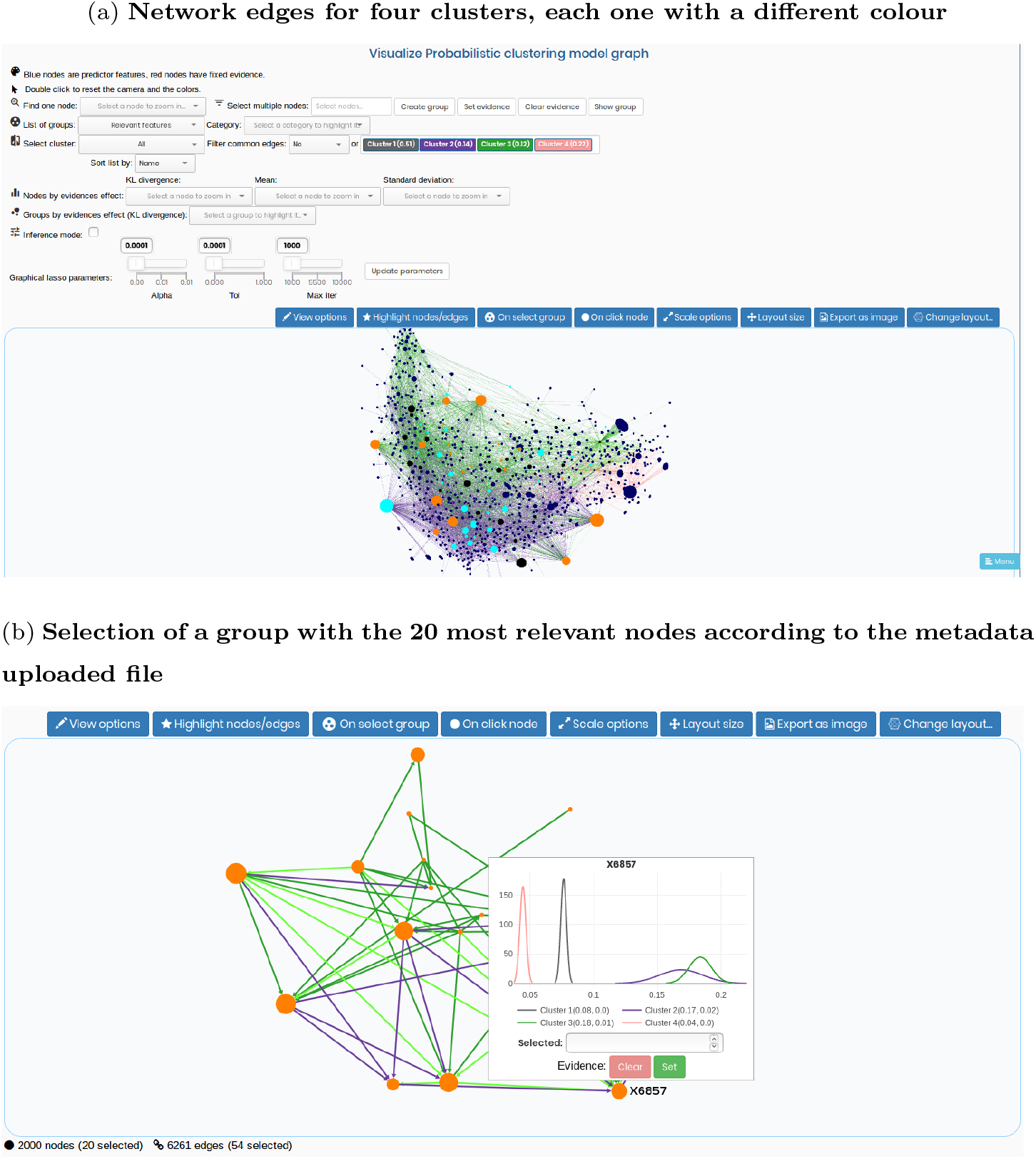
BN-based probabilistic clustering model of 2000 nodes of the human brain genome. (a) The upper part of the image also presents the cluster weights. Node sizes are adjusted to highlight the most important nodes with the betweenness centrality algorithm. Nodes colours are according to external metadata to organize them in three groups given their importance. (b) The plot displays the parameters of gene X6857. Each of the four clusters (different colours), presents a Gaussian distribution. In this example, we can easily notice that the most probable cluster assignation for this gene is cluster 1 (in grey), *p*(*X*6857|*C* = *c*_1_).

Finally, we express the joint probability distribution of ***X*** (Equation 3) factorized according to Equation 4. We call this BN-based probabilistic clustering [76],

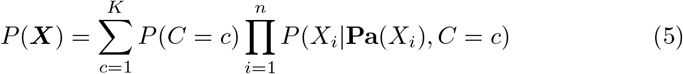

Therefore, inference can be performed on each graph corresponding to a cluster without affecting the other cluster CPDs. For instance, we can fix the evidence for the distribution of a gene, as *X_i_ = e*, given a cluster *C = c*, where *e* is a scalar value, and then query another gene to determine how its CPD for that cluster has changed, *P*(*X_j_*|*C = c, X_i_ = e*).

The obtained BN can be exported as an SVG image or as a CSV file containing the graph information about the arcs between the nodes. This exported file can be loaded subsequently in another session to continue working. Finally, it is important to acknowledge that the user data in a session remains in our servers for 48 h since the last modification of the data. This limit is imposed by our hardware limitations. To overcome this limitation, a user can always create new sessions, and the data will be stored again for 48 h. Users are also encouraged to deploy their own server instance to modify the framework according to their needs.

## 4. Discussion

Here, we review future directions for BayeSuites by first introducing new use cases for which we believe this tool could be of great interest, and then, we also indicate potential extensions and new functionalities that would make BayeSuites even more complete.

We believe that the ease of use will be helpful in initiating collaborations between experts of multiple disciplines. This will be extremely important for the adoption of these models by experts of other disciplines who are not used to programming or software engineering, such as some neuroscientists or physicians.

A useful use case could be the use of a private server instance in closed local networks environments, such as hospitals, clinical laboratories, or companies. A workflow could be easily designed to have a clear pipeline to process the data with machine learning techniques. New data sources connections could be implemented to automatically plug into the data acquisition machines. In addition, some type of specific pre-processing for the data could be implemented in NeuroSuites (e.g., for genomic data it could be the removal of irrelevant genes and the inclusion of domain knowledge about the most important genes). Further, the experts could analyse the data with the BayeSuites framework. The web characteristics of the frameworks would make the tool available in a web browser for each employee in the local network without the need of installing the software on their computer.

Finally, we also believe this simplicity could be a great aid for educational purposes when teaching BNs allowing the theorical properties to be shown in a dynamic environment.

The framework aims to be a complete product; however, this is an extremely large research field, and at the time of writing this paper it does not include all the existing state-of-the-art functionalities. Its extensibility properties can make it possible to include numerous extensions and implement new functionalities.

A useful implementation to be included would be some inference algorithms for discrete BNs. We have provided the support to learn and visualize discrete parameters in BNs. However, we have not included yet any inference algorithm for them owing to the development time constraints and the difficulty to visualize the changes in the parameters when there are many parameters per node and numerous nodes. Moreover, massive datasets in various neuroscience fields, such as genomics and electrophysiology, comprise only continuous features.

Another interesting extension would be the inclusion of dynamic BNs [77]. The steps to implement this would be similar to the ones described in the last section to include BN-based clustering models. However, there would be an increased complexity to visualize the network for each timeframe and for performing new types of inferences (e.g., filtering, smoothing, etc.).

Finally, we want to highlight that NeuroSuites also offers different tools for other neuroscience domains, such as morphological reconstructions and microscopy data visualization. However, although this framework is designed focusing on the neuroscience field, many other tools can also be used in other research fields. Developers can modify the platform to target a different research field. However, it is also important to note that no modifications are needed if the user wants to upload his own dataset and learn a probabilistic graphical model and interpret it, despite the neuroscience background theme of the website. For instance, the use case that we followed here needs a specific BN structure learning algorithm designed for genomics (FGES-Merge) along with all the visualization tools for understanding its massive network. However, for other domains, where datasets are relatively smaller, other algorithms could also be applied.

## 5. Conclusion

In this paper, we have presented BayeSuites, which aims not only to get the best from the proprietary and open source worlds but also to extend it. The result is an open source web framework for learning, visualizing, and interpreting BNs, with the peculiarity of being the first web framework that is scalable and able to manage massive BNs of tens of thousands of nodes and edges. This ability is required to overcome the main obstacles for managing massive BNs, and accurate and fast methods for structure learning, visualization and inference were developed. This development was done by providing a friendly and interactive user experience.

To test our tool, we extensively compared BayeSuites with the major BN tools currently available; we divided the necessary BNs functionalities into four categories: scalability, extensibility, interoperability and ease of use. We conclude that, presently, BayeSuites is the only tool that fully incorporates all these functionalities for massive BNs.

Finally, we showcased the utility of BayeSuites by providing two real use cases of the entire process of learning, visualizing and interpreting BNs from genomic data obtained from the Allen Institute for Brain Science.

## 6. Data availability

Our production server on https://neurosuites.com/morpho/ml_bayesian_network can be freely accessed, where all the futures updates will be live. We also provide access to the NeuroSuites source code repository in https://gitlab.com/mmichiels/neurosuite. The BNs used in the examples for showcasing BayeSuites can be found in https://gitlab.com/mmichiels/fges_parallel_production/tree/master/BNs_results_paper

## 7. Author contributions

Mario Michiels designed the software architecture, developed the software and wrote the manuscript. Pedro Larrañaga and Concha Bielza conceived the project, oversaw the development process, contributing with new ideas and corrections, and reviewed the manuscript. All authors gave final approval for publication and agree to be held accountable for the work performed therein.

## 10. Acknowledgments

The authors would like to thank Sergio Paniego for his help in the development of the BayeSuites visualization tool, Nikolas Bernaola for his assistance in programming the continuous inference engine for BNs, and Fernando Rodriguez-Sanchez for his research in BN-based probabilistic clustering models and his help in reviewing that section in this paper.

